# Transcriptional repression of SOX2 by p53 in cancer cells regulates cell identity and migration

**DOI:** 10.1101/2024.11.14.623640

**Authors:** Patricia Lado-Fernández, Jéssica M Vilas, Tânia Fernandes, Carmen Carneiro, Sabela Da Silva-Álvarez, Valentín Estévez-Souto, Pablo Pedrosa, Miguel González-Barcia, Luis E. Abatti, Jennifer A. Mitchell, Carmen Rivas, Gema Moreno-Bueno, Anxo Vidal, Manuel Collado

## Abstract

During cancer development and progression, many genetic alterations lead to the acquisition of novel features that confer selective advantage to cancer cells and that resemble developmental programs. SOX2 is one of the key pluripotency transcription factors, expressed during embryonic development and active in adult stem cells. In cancer, *SOX2* is frequently dysregulated and associated with tumor stemness and poor patient survival. *SOX2* expression is suppressed in differentiated cells by tumor suppressor proteins that form a transcriptional repressive complex. We previously identified some of these proteins and found that their absence combined with deficiency in Trp53, leads to maximal dysregulated expression of *Sox2*. Using cancer cell lines of different origin and with different p53 status, we show here that manipulating *TP53* to restore or decrease its activity results in repression or induction of *SOX2*, respectively. Mechanistically, we observed that the regulation of *SOX2* expression by *TP53* is transcriptional and identified Trp53 bound to the promoter region and the *SRR2* enhancer of *Sox2*. Forcing high levels of *SOX2* in cancer cells leads to morphological changes that molecularly correspond to the acquisition of a more mesenchymal phenotype correlating with an increased migratory capacity. Finally, the analysis of human breast cancer samples shows that this correlation between TP53 status, levels of expression of SOX2 and a more metastatic phenotype is also observed in cancer patients. Our results support the notion that lack of TP53 in tumor cells results in deregulated expression of developmental gene *SOX2* with phenotypic consequences related with increased malignization.

**Graphical Abstract:** 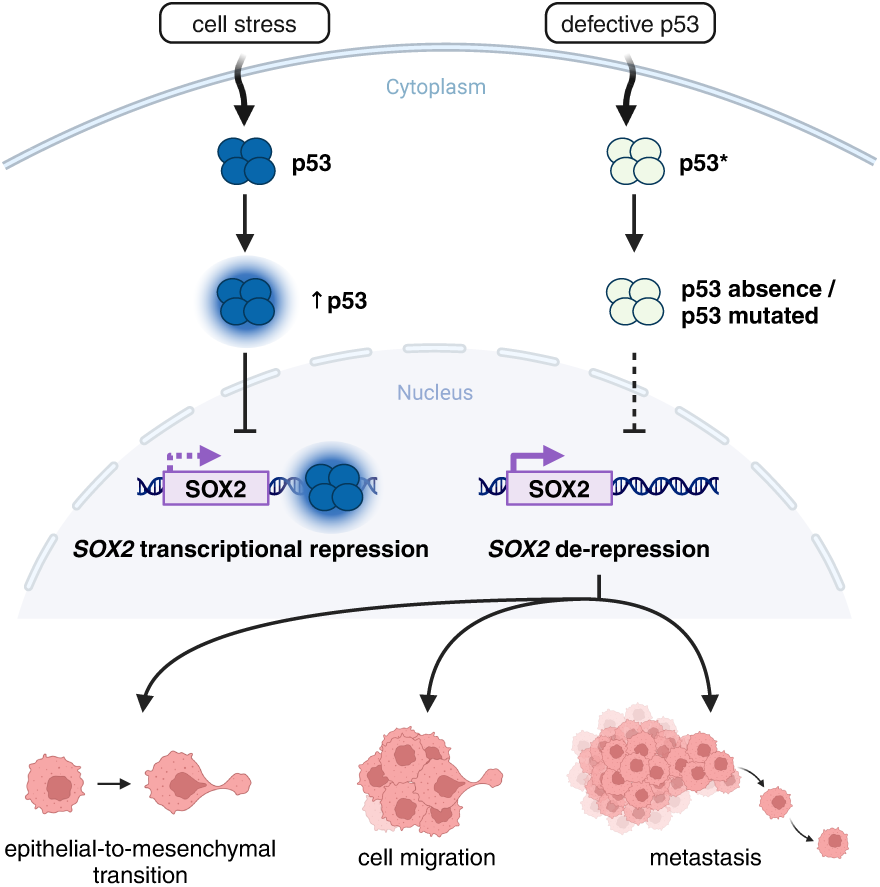

**Novelty and Impact:** Our work demonstrates that TP53 regulates transcriptionally *SOX2* through binding to its promoter and the *SRR2* enhancer. Increased expression of *SOX2* promotes an EMT and increased migration. Breast cancer patients with genetic alteration of TP53 show increased levels of SOX2 and a more metastatic phenotype.

This work is a key demonstration of an additional effect of genetic alterations of *TP53*, leading to de-repression of *SOX2* and alteration of cancer cell plasticity with important phenotypic consequences.

## INTRODUCTION

Cancer progression is driven by genetic mutations in genes that confer a selective advantage to tumor cells. Along tumor development, cancer cells acquire remarkable biological properties that allow them to sustain robust proliferation, escape tumor suppressor control, resist cell death, replicate indefinitely, induce angiogenesis, and ultimately migrate and invade distant organs ^1^. Some of these properties are reminiscent of features that are typical of embryonic development, and the reactivation of developmental programs has been identified during tumor progression. Alternatively, these new malignant attributes could emerge from the acquisition of adult stem/progenitor features by cancer cells.

SRY-box transcription factor 2 (*SOX2*) is an essential gene to maintain pluripotency in early embryonic cells of the inner cell mass of the blastocyst, and it is a key factor to reprogram differentiated adult somatic cells back into a pluripotent stem cell stage ^2–4^. Its expression can be found in progenitors of several epithelia during fetal development while, after birth, small populations of Sox2 expressing cells seem to play crucial roles regulating endodermal and ectodermal tissue regeneration and homeostasis ^5^.

The role of Sox2 has also been investigated in the context of cancer. *SOX2* expression is high in tumors of different origin, and its gene amplification has even been identified as driving lung and esophageal squamous cell carcinomas ^6,7^. Several studies indicate that *SOX2* overexpression provides tumor cells with advantageous properties for tumor development, including increased cellular proliferation, invasion, migration, anchorage-independent growth, evasion of apoptosis and drug resistance, while *SOX2* knockdown has been linked to reduced tumor progression and decreased resistance to cancer treatments ^8,9^.

Little is known however, regarding the regulation of Sox2 in cancer. In previous work from our group, while investigating the transcriptional regulation of *SOX2* by CDK inhibitor p27Kip1 ^10^ and the retinoblastoma family of pocket proteins ^11^, we realized that a low-level expression of *Sox2*, causing dramatic phenotypic consequences, was sufficient to activate its own endogenous transcription provided that *Trp53* was downregulated. These results pointed to a potential novel function for Trp53 repressing the expression of *Sox2* in differentiated cells and a novel mechanism for deregulated *Sox2* expression in cancer.

To further investigate this potential link between the tumor suppressor activity of p53 and de-regulation of *SOX2* in cancer, we decided to manipulate *TP53* levels in cancer cells that express wild type *TP53* or that lack expression of this tumor suppressor. We found that enforced expression of exogenous *TP53*, or DNA damage induction of the endogenous gene, leads to a transcriptional repression of *SOX2*. This regulation seems to be mediated, at least in part, by binding of TP53 to the *SOX2* promoter and to its proximal enhancer *SRR2* (*Sox2* Regulatory Region 2). Conversely, downregulation of endogenous *TP53* seems to alleviate *SOX2* repression leading to an increased expression of *SOX2*. Phenotypically, we found that cancer cells with higher level expression of *SOX2* undergo a morphological change accompanied by expression of markers and mediators of the epithelial-to-mesenchymal transition. Functionally, this altered phenotype caused by increased levels of SOX2 correlated with an increase in the migratory capacity of cancer cells, suggesting a link between high level expression of *SOX2* in cancer and increased migration, and possibly a more metastatic capacity. This hypothesis was further supported by data from breast cancer patient derived samples that showed a positive correlation between higher expression of *SOX2*, mutation in *TP53*, and a more metastatic behavior.

## MATERIALS AND METHODS

### Cell line generation

NCI-H1299 (RRID:CVCL_0060), U-373MG ATCC (RRID:CVCL_2219), T-47D (RRID:CVCL_0553), A-549 (RRID:CVCL_0023), U-87MG ATCC (RRID:CVCL_0022), MCF-7 (RRID:CVCL_0031) and HEK293T (RRID:CVCL_0063) cell lines were obtained from the ATCC. T653 and T731 cell lines were a kind gift from José Antonio Costoya (CiMUS, USC). All cell lines were cultured in DMEM (Dulbecco’s Modified Eagle Medium) (Sigma-Aldrich) supplemented with 10% FBS (fetal bovine serum) (Sigma-Aldrich), 1% glutamine (Sigma-Aldrich) and 1% penicillin/streptomycin (Sigma-Aldrich); maintained in an incubator at 37° C and 5% CO_2_ atmosphere. Human cell lines have been authenticated using STR profiling and all cell lines were routinely tested for mycoplasma.

To generate TP53-overexpressing cells, target cells were transfected with expression plasmid pcDNA3-p53 (a gift from R. T. Hay) ^12^, and pcDNA3 (Invitrogen) as a control, in the presence of 1 mg/mL polyethylenimine reagent (PEI) (Polysciences). To generate TP53-knockdown cell lines, human cancer cells were transduced with pRetroSuper-Blast-shp53 and murine cancer cells were transduced with pRetroSuper-puro-shp53, or with the respective controls pRetroSuper-Blast and pRetroSuper-puro (all of them a kind gift from Reuven Agami).

To generate cell lines with increased endogenous *SOX2* levels, cells were transduced with PL-SIN-EOS-C(3+)-EiP (a gift from James Ellis; Addgene #21313) and selected with 1 µg/mL of puromycin (Millipore) ^13^. *SOX2*-overexpressing cell lines were generated by retro or lentiviral transduction with pMXs-Sox2 (a gift from Shinya Yamanaka; Addgene #13367) ^4^, and pBabe-puro-IRES-EGFP (a gift from L. Miguel Martins; Addgene #14430) as a control, or pLV-tetO-Sox2 (a gift from Konrad Hochedlinger; Addgene #19765) ^14^ and the transactivator FUW-M2rtTA (a gift from Rudolf Jaenisch; Addgene #20342) ^15^, followed by treatment with 0.5 µg/mL of doxycycline.

For retroviral and lentiviral transduction, HEK293T cells were co-transfected with the plasmid of interest and the corresponding packaging plasmids in equal proportions, in the presence of 1 mg/mL PEI reagent. Third-generation lentiviral packaging plasmids were pLP1, pLP2, pLP-VSVG (ViraPower Lentiviral Packaging Mix, K497500, Invitrogen); second-generation lentiviral packaging plasmids were psPAX2 (a gift from Didier Trono; Addgene #12260) and pLP-VSVG and retroviral packaging plasmids were pCL-Eco (a gift from Inder Verma; Addgene #12371) ^16^ and pLP-VSVG. 36 hours after transfection, we collected viral supernatants every 12 hours for three consecutive rounds. The supernatants were filtered through a 0.45 μm filter, supplemented with 8 μg/mL of polybrene (Sigma-Aldrich) and added to target cells. Cell cultures transduced with plasmids with antibiotic resistance genes were selected with 1 µg/mL puromycin (Millipore) or 4-10 µg/mL blasticidin (Sigma-Aldrich).

### RT-qPCR

To measure gene expression, total RNA was extracted using the NucleoSpin RNA Plus kit (Macherey-Nagel) following the manufacturer’s protocol, and retrotranscribed into cDNA using the High-Capacity cDNA Reverse Transcription Kit (Applied Biosystems) according to the manufacturer’s protocol. Quantitative Real Time PCR (RT-qPCR) was performed using the NZYSpeedy qPCR Green Master Mix reagent (2X) ROX (NZYTech) and the QuantStudio 3 Real-Time PCR System (Applied Biosystems). Each reaction was performed in triplicate and consisted of 33 ng of cDNA, 0.25 μM primers, 5 μL of Master Mix reagent in a final volume of 10 μL per reaction. RNA expression values were relativized to the expression levels of the housekeeping gene *GAPDH*. The results were analyzed with the QuantStudio 3 Real-Time PCR Software (Applied Biosystems). Primer sequences were as follows:

*GAPDH*: 5’-TCCATGACAACTTTGGCATCGTGG-3’, 5’-GTTGCTGTTGAAGTCACAGGAGAC-3’;

*TP53*: 5’-CCGCAGTCAGATCCTAGCG-3’, 5’-AATCATCCATTGCTTGGGACG-3’;

*CDKN1A*: 5’-CCTGTCACTGTCTTGTACCCT-3’, 5’-GCGTTTGGAGTGGTAGAAATCT-3’;

*SOX2*: 5’-GGGAAATGGGAGGGGTGCAAAAGAGG-3’, 5’-TTGCGTGAGTGTGGATGGGATTGGTG-3’;

*Cdkn1a*: 5’-GTGGGTCTGACTCCAGCCC-3’, 5’-CCTTCTCGTGAGACGCTTAC-3’;

*Sox2*: 5’-TAGAGCTAGACTCCGGGCGATGA-3’, 5’-TTGCCTTAAACAAGACCACGAAA-3’;

*Trp53*: 5’-TGAAACGCCGACCTATCCTTA-3’, 5’-GGCACAAACACGAACCTCAAA-3’;

*CDH1*: 5’-CGAGAGCTACACGTTCACGG-3’, 5’-GGGTGTCGAGGGAAAAATAGG-3’;

*CDH2*: 5’-AGCCAACCTTAACTGAGGAGT-3’, 5’-GGCAAGTTGATTGGAGGGATG-3’;

*SNAI1*: 5’-TCGGAAGCCTAACTACAGCGA-3’, 5’-AGATGAGCATTGGCAGCGAG-3’ and

*TWIST1*: 5’-GTCCGCAGTCTTACGAGGAG-3’, 5’-GCTTGAGGGTCTGAATCTTGCT-3’.

### Western blot

To analyze protein expression, cell extracts were resuspended in RIPA buffer (150 mM NaCl, 10 mM Tris-HCl pH 7.5, 0.1% sodium dodecyl sulfate, 1% Triton X100, 5 mM EDTA pH 8.0, 1% deoxycholate and sodium salt) containing protease inhibitors (1 mM sodium orthovanadate; 1 mM PMSF; 1 mM DTT; 4 mM NaF; protein inhibitor cocktail, P8340 Sigma-Aldrich). Protein concentration was quantified with the DC Protein Assay Kit (Bio-Rad) and 25 μg of total cellular protein were electrophoresed in 12% polyacrylamide gels. After transferring to a PVDF membrane, the blot was blocked with 5% milk solution in TTBS (20 mM Tris-HCl pH 7.5, 150 mM NaCl, 0.05% Tween-20). Membranes were incubated at 4 °C overnight with primary antibodies (TP53: sc-6243, Santa Cruz Biotechnology, 1:500; P21: sc-6246, Santa Cruz Biotechnology, 1:500; SOX2: #2748, Cell Signaling, 1:1000; β-TUBULIN: #2146, Cell Signaling, 1:1000; GAPDH: sc-32233, Santa Cruz Biotechnology, 1:2000; β-ACTIN: sc-8432, Santa Cruz Biotechnology, 1:1000). Secondary antibodies (anti-mouse IgG-HRP: #31430, Invitrogen, 1:10,000; anti-rabbit IgG-HRP: #31460, Invitrogen, 1:10,000) were incubated for 1 h. Chemiluminescence detection was performed using SuperSignal West Pico PLUS Chemiluminescent Substrate (Thermo Scientific) in a ChemiDoc MP Imaging System (BIO-RAD).

### Luciferase reporters

To analyze the activity of regulatory regions, cells were co-transfected with luciferase reporter plasmids PG13-luc (wt p53 binding sites) (a gift from Bert Vogelstein; Addgene #16442) ^17(p1)^ and pGL3-Sox2 luc (a gift from Richard G. Pestell) in presence of polyethylenimine. After 48 hours, the cultures were lysed with lysis buffer (Roche) and luciferase assay buffer (25 mM glycylglycylglycine pH 7.8, 15 mM phosphate buffer pH 7.8 (KH_2_PO_4_ and K_2_HPO_4_), 15 mM MgSO_4_, 4 mM EGTA, 2 mM ATP, 1 mM DTT). Luciferase activity was measured using the Mithras LB 940 luminometer (Berthold Technologies) after adding D-luciferin (Sigma-Aldrich) with the automatic injector. To ensure transfection efficiency, cells were co-transfected with the plasmid pBP-LacZ (a gift from Paul Freemont; Addgene #72948) ^18^ and β-galactosidase activity was measured. Cell extracts were added to Z-buffer (40 mM NaH_2_PO_4_, 60 mM Na_2_HPO_4_, 10 mM KCl, 1 mM MgSO_4_, β-mercaptoethanol and 2-nitrophenyl) for this purpose. Β-D-galactopyranoside (Sigma-Aldrich) was incubated at 37° C for 8 hours. The optical density was measured at 450 nm using Mithras LB 940 (Berthold Technologies).

### Chromatin immunoprecipitation

Chromatin immunoprecipitation (ChIP) was performed as described in ^19^, with minor adjustments. After chromatin extraction, lysates were sonicated at 20% amplitude, 15 seconds ON, 30 seconds OFF for 7 minutes using a Branson Sonifier 550 (Branson Ultrasonics). Antibodies used for ChIP were anti-goat IgG (G4018, Sigma-Aldrich) and p53 (FL-393) (sc-6243, Santa Cruz Biotechnology). DNA samples were purified with UltraPure Phenol:Chloroform:Isoamyl Alcohol (25:24:1, v/v) (15593-031, Invitrogen) and eluted in 80 µL H_2_O. Quantitative Real Time PCR was performed in triplicates and each reaction consisted of 2 µL sample, 0.25 μM primers and 5 μL of NZYSpeedy qPCR Green Master Mix reagent (2X) ROX (NZYTech) in a final volume of 10 μL per reaction, using the QuantStudio 3 Real-Time PCR System (Applied Biosystems). Primer sequences were following:

*Cdkn1a promoter*: 5’-CCAGAGGATACCTTGCAAGGC-3’, 5’-TCTCTGTCTCCATTCATGCTCCTCC-3’;

*Sox2 promoter*: 5’-CCTAGGAAAAGGCTGGGAAC-3’, 5’-GTGGTGTGCCATTGTTTCTG-3’;

*Sox2-SRR1*: 5’-TCCCCCAATACTGGTGGTCGTCA-3’, 5’-GAAGGCGAACGGCAGGGGAC-3’;

*Sox2-SRR2*^1^: 5’-ATTTATTCAGTTCCCAGTCCAAGC-3’, 5’-CCCTCTCCCCCCACGC-3’ and

*Sox2-SRR2*^2^: 5’-CGTGGTAATGAGCACAGTCG-3’, 5’-AGGCTGAGTCGGGTCAATTA-3’.

### Proliferation and clonogenicity assays

To analyze cell proliferation, A-549 and MCF-7 cells were seeded at a density of 8×10^3^ and 1×10^4^ cells, respectively, in 24-well plates in triplicate for each condition. Cells were counted every 2 days using the LUNA II Automated Cell Counter (Logos Biosystems). For the clonogenicity assay, 2×10^3^ A-549 or MCF-7 cells were seeded in 6-well plates, in triplicate for each condition. After 12-14 days cell colonies were fixed with 4% paraformaldehyde (PFA) and stained with 0.05% crystal violet solution. Images were obtained after scanning the plates with the Epson Perfection V550 Photo scanner. For quantification, the staining was eluted with 10% acetic acid solution and the optical density was measured at 570 nm in the spectrophotometer Epoch 2 (Biotek).

### Migration assay

Cell migration capacity was assessed by migration through 8 µm *Transwell* inserts (Falcon) in 24-well plates. A-549 and MCF-7 cells were seeded in the upper chamber at a density of 2.5×10^4^ cells in serum-free medium, in triplicate. In the lower chamber, the medium was supplemented with 10% FBS to promote cell migration. After 16 hours for A-549 or 48 hours for MCF-7, cells were fixed with 4% PFA, permeabilized with 100% methanol, and stained with 0.05% crystal violet solution. After removing the cells in the upper chamber, we took images with Axio Vert.A1 Microscope (Zeiss) and the number of migrated cells was quantified using ImageJ.

### Immunohistochemistry

This study utilized TMA (tissue microarray) and breast carcinoma samples obtained from a previously described cohort ^20^. Tumor sections, two micrometers thick, were deparaffinized using BOND Dewax Solution (Leica Biosystems) and subsequently rehydrated with a series of ethanol solutions. Immunohistochemistry was performed using a BOND RXm autostainer (Leica Biosystems), employing BOND Epitope Retrieval Solutions and the BOND Polymer Refine Detection kit (Leica Biosystems) following the manufacturer’s protocols. Images were captured using Aperio eSlide Manager software (Leica Biosystems). SOX2 and TP53 levels were categorized according to ^21^.

### Statistical analysis

All data are represented as mean ± standard deviation. Statistical significance was calculated using the two-tailed Student’s t-test for most experiments, except for immunohistochemistry experiments, which were analyzed using the two-tailed Fisher’s exact test. P-values less than 0.05 were considered statistically significant: *** *p*<0.001; ** *p*<0.01; * *p*<0.05; n.s. not significant.

## RESULTS

### Tumor suppressor *TP53* regulates *SOX2*

To explore the contribution of tumor suppressor TP53 to the regulation of the expression of SOX2 in cancer cells, we selected lung (NCI-H1299 and A-549), glioma (U-373MG and U-87MG), and breast (T-47D and MCF-7) tumor cells lines with defective or wild type TP53, respectively; and transfected them with a plasmid expressing TP53 or an short hairpin RNA (shRNA) against TP53 to restore or decrease the expression of the tumor suppressor, accordingly. When we transfected *TP53* deficient NCI-H1299, U-373MG and T-47D cells with a plasmid coding for TP53, we observed different levels of expression of the tumor suppressor protein and of its target gene, *CDKN1A* (coding for p21). Following the increased expression of TP53 in all the tumor cell lines we observed a reduction of SOX2, both at mRNA (Fig. 1A) and protein levels (Fig. 1B). Conversely, when we introduced a shRNA targeting *TP53* in wild type A-549, U-87MG and MCF-7 cells, the mRNA (Fig. 1C) and protein levels (Fig. 1D) of *TP53* and *CDKN1A* (and of its encoded protein, p21) were reduced, as expected, while at the same time SOX2 was increased (Fig. 1C and 1D).

**Fig. 1.**
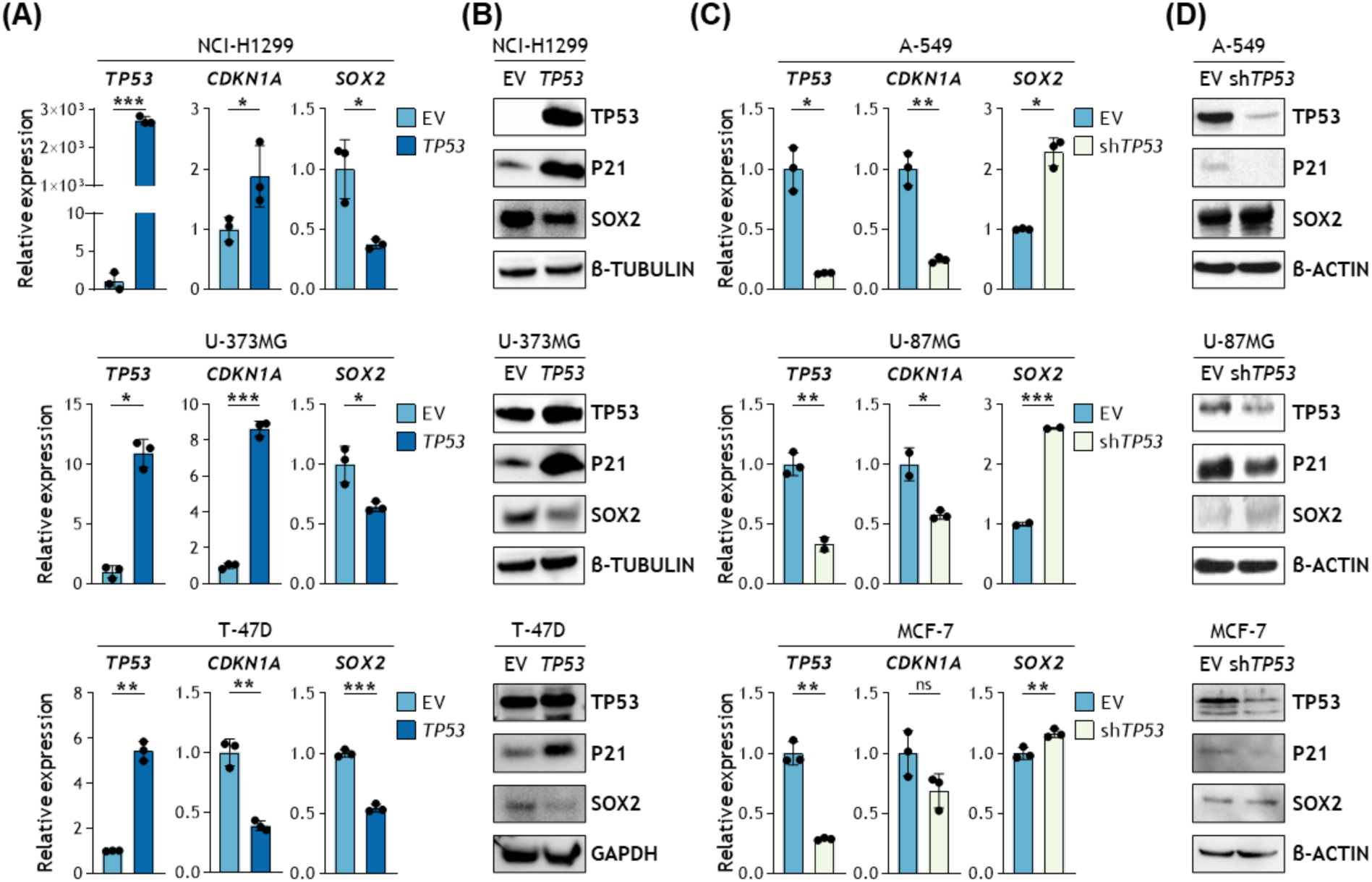
Effects of altered TP53 expression on SOX2 levels. **(A)**, *TP53* (left), *CDKN1A* (middle) and *SOX2* (right) relative mRNA levels by RT-qPCR in *TP53* defective NCI-H1299, U-373MG and T-47D tumor cell lines overexpressing *TP53*. **(B)**, TP53, P21 and SOX2 protein levels by western blot in *TP53* defective NCI-H1299, U-373MG and T-47D tumor cell lines overexpressing *TP53*. **(C)**, *TP53* (left), *CDKN1A* (middle) and *SOX2* (right) relative mRNA levels by RT-qPCR in *TP53* wild type A-549, U-87MG and MCF-7 tumor cell lines with silenced *TP53* expression. **(D)**, TP53, CDKN1A and SOX2 protein levels by western blot in *TP53* wild type A-549, U-87MG and MCF-7 tumor cell lines with silenced *TP53* expression.

In order to corroborate these findings without recurring to genetic manipulation, we decided to investigate the effect of using a chemotherapeutic drug that causes DNA damage and consequently activates TP53, namely doxorubicin, on the expression of SOX2. For this, we used two tumor cell lines derived from gliomas originated in transgenic K*ras*G12V mice, T653 and T731. Addition of doxorubicin (0.5 μg/mL) to these cells caused a time dependent increase in Trp53 protein that was accompanied by an increase of its target, p21, and a concomitant decrease in the levels of Sox2 protein in both cell lines (Fig. 2A). Again, the analysis of *Sox2* mRNA revealed that these reduced levels of Sox2 were caused by a downregulation of its mRNA after activation of Trp53 and of *Cdkn1a* by doxorubicin (Fig. 2B).

**Fig. 2.**
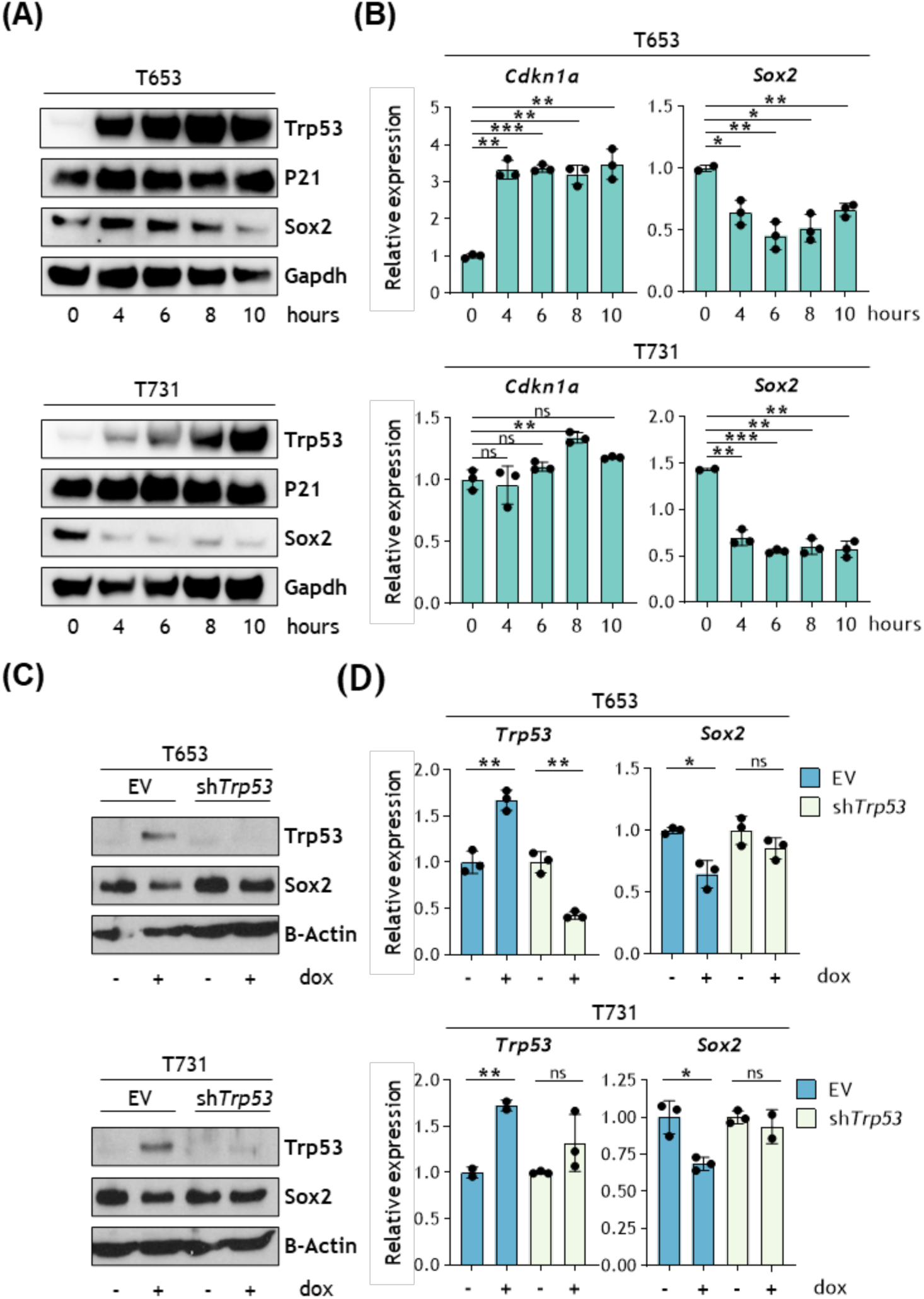
The activation of *Trp53* by DNA damage decreases *Sox2* levels. **(A)**, Trp53, P21 and Sox2 protein levels assessed by western blot in T653 (top) and T731 (bottom) mouse glioma cell lines treated with 0.5 µg/ml doxorubicin at different times. **(B)**, Relative mRNA levels by RT-qPCR of the Trp53 target gene *Cdkn1a* and *Sox2* in T653 and T731 cells treated with 0.5 µg/ml doxorubicin at different times. **(C)**, Trp53, P21 and Sox2 protein levels by western blot in T653 and T731 cells previously transduced with an shTrp53 and treated with 0.5 µg/ml doxorubicin for 10 hours. **(D)**, Relative mRNA levels by RT-qPCR of *Trp53* and *Sox2* in T653 and T731 cells previously transduced with an shTrp53 and treated with 0.5 µg/ml doxorubicin for 10 hours. doxo, doxorrubicin.

To prove that this reduction in *Sox2* mRNA and protein after doxorubicin treatment of the glioma cells is due to the activation of *Trp53* and not the result of an off-target effect of doxorubicin, we repeated the experiment adding the drug to cells previously transduced with an shRNA construct targeting *Trp53*. Again, we observed that doxorubicin caused a sharp increase in the levels of Trp53, as expected, and this change correlated with a reduction in the levels of *Sox2*, both at the protein and mRNA levels (Fig. 2C and 2D, respectively). Interestingly, introduction of the shRNA against *Trp53* impeded the increase in *Trp53* after doxorubicin addition and this in turn prevented the repression of *Sox2* expression (Fig. 2C and 2D).

These results clearly show that increased genetic expression of *TP53* or its induction after DNA damage in cancer cells results in downregulation of SOX2 expression and that, conversely, inactivation of *TP53* in cancer cells retaining a wild type tumor suppressor protein alleviates SOX2 transcriptional repression.

### The regulation of *SOX2* by tumor suppressor *TP53* is transcriptional

To address if the effect of *TP53* on *SOX2* expression was produced at the transcriptional level, we transfected lung and glioma tumor cell lines carrying wild type or defective *TP53*, with reporter constructs expressing luciferase under the control of a promoter containing 13 copies of the Tp53-binding consensus sequence (pG13-luc), as a positive control for Tp53 activity, or with a reporter expressing luciferase under the control of *SOX2* promoter (pGl3-Sox2-luc). Using this system, we observed that overexpression of *TP53* caused an increase in luciferase activity in the positive control pG13-luc cells, as expected, and a decrease in luciferase from the *SOX2* promoter reporter plasmid pGl3-Sox2-luc (Fig. 3A, upper panels). Conversely, knockdown of *TP53* reduced the expression of luciferase from the control plasmid, pG13-luc, and stimulated luciferase expression from *SOX2* promoter, confirming that *TP53* expression negatively affects the transcription of *SOX2* (Fig. 3B, lower panels).

**Fig. 3.**
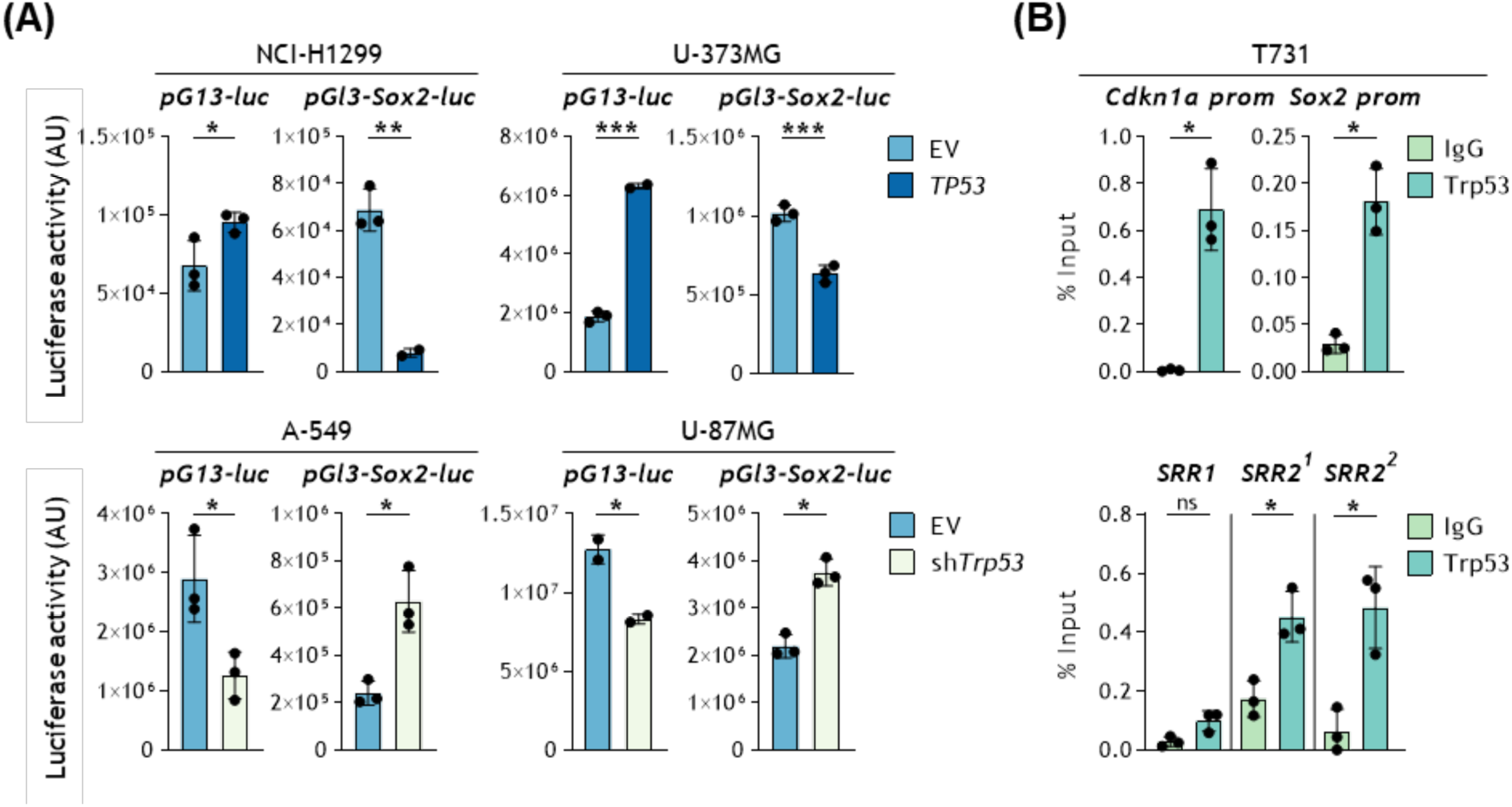
The alteration of TP53 levels modulates *SOX2* activity transcriptionally in lung cancer and glioma cell lines. **(A)**, Luciferase activity from the *CDKN1A* promoter (pG13-luc) or the *SOX2* promoter (pGl3-Sox2-luc) in *TP53* defective tumor cell lines NCI-H1299 and U-373MG overexpressing *TP53* (top) and in *TP53* wild type cell lines A-549 and U-87MG (bottom). **(B)**, Analysis of TP53 binding by Trp53 induction with 0.5 µg/ml doxorubicin for 10 hours followed by chromatin immunoprecipitation with a Trp53 antibody and RT-qPCR of the *Cdkn1a* promoter, the *Sox2* promoter, the *SRR1* and the *SRR2* (with 2 different primer pairs).

Next, we wondered if we could identify Trp53 bound to regulatory regions on the *Sox2* locus. For this, we performed chromatin immunoprecipitation followed by RT-qPCR with primers amplifying the promoter region or two previously described enhancers downstream of *Sox2* open reading frame, namely the *SRR1* and the *SRR2*. ChIP using specific anti-Trp53 antibodies allowed the detection of the *Cdkn1a* promoter, a well-known transcriptional target of Trp53, in T731 cells after doxorubicin treatment. Similarly, ChIP after DNA damage led to the detection of *Sox2* promoter as well as of the *SRR2* using two different pairs of primers for this enhancer region, but not of the *SRR1* enhancer (Fig. 3B).

Altogether, our results clearly show that TP53 represses *SOX2* gene expression in cancer cells acting at the transcriptional level, at least through binding to the *SOX2* promoter and its *SRR2* enhancer.

### Increased levels of SOX2 in cancer cells do not alter proliferation

Next, we wondered what the effect would be of expressing higher levels of SOX2 in a cancer cell. To address this question, we set up three different systems to achieve higher gradual expression levels of Sox2. For the first one, we took advantage of a SOX2 reporter plasmid carrying tandem repeats of an enhancer element containing *Sox2*/*Oct4* binding sites upstream of a promoter derived from a murine early transposon active in embryonic stem cells that drives the expression of GFP (Green Fluorescent Protein) and puromycin resistance (EOS-GFP). Introduction of this construct in A-549 and MCF-7 cells renders a small fraction of these cell cultures positive for GFP and resistant to puromycin (Fig. S1A and S1B). Antibiotic selection of this population enriches for tumor cells showing the highest natural level of expression of SOX2 within the cancer cell line, as we confirmed by Western blot and RT-qPCR (Fig. S1C). As a second approach, we used retroviral expression of Sox2 using pMXs-Sox2-IP. After transduction and selection of the cell population carrying the plasmid, we confirmed the elevated levels of Sox2 in these cells by Western blot and RT-qPCR (Fig. S1D). Finally, we used a tetracycline-inducible system to overexpress Sox2, pLV-tetO-Sox2, showing the highest level of expression of Sox2 protein and mRNA of the three systems (Fig. S1E).

Using these cell models with different degrees of expression of SOX2, we observed that cells expressing higher levels of SOX2 did not show increased proliferation when followed for 8 days in culture (Fig. S2A). On the contrary, when Sox2 expression was maximal cells appeared to have an impaired proliferative rate (Fig. S2A, bottom panels). Moreover, when we tested the capacity of SOX2 overexpressing tumor cells to form colonies at low density after 14 days in culture, the results indicated that SOX2 expression does not confer a proliferative advantage to tumor cells (Fig. S2B) or, if anything, it represents an obstacle for efficient proliferation when SOX2 is very highly expressed (Fig. S2B, bottom panels).

### Increased levels of SOX2 in cancer cells induce features of epithelial-to-mesenchymal transition

The microscopic inspection of these cell cultures revealed however morphological changes in the cells expressing higher levels of SOX2, with cells showing a more elongated fusiform shape, resembling a more mesenchymal phenotype in the epithelial cancer cell lines A-549 (Fig. S3A, S3C and S3E) and MCF-7 (Fig. S3B, S3D and S3F), and with a stronger effect caused by the two overexpression systems, pMXs-Sox2-IP (Fig. S3C and S3D) and pLV-tetO-Sox2 (Fig. S3E and S3F). This prompted us to evaluate the possibility of an induced epithelial-to-mesenchymal transition (EMT) caused by the forced expression of SOX2 in A-549 and MCF-7. For this, we first checked the expression of a classical epithelial marker, *CDH1* (coding for E-Cadherin), and a mesenchymal marker, *CDH2* (coding for N-Cadherin), using the three systems of increased expression of SOX2. We confirmed that, for the most part, the levels of *CDH1* were reduced while for *CDH2* they were increased when SOX2 was overexpressed (Fig. 4A and 4B), confirming the induction of a change in the phenotype of the epithelial cancer cell lines A-549 and MCF-7 to a more mesenchymal one. Apart from the expression of cell identity markers we decided to check the expression of some of the most commonly described mediators of the process of EMT, transcription factors *SNAIL1* and *TWIST1*. Both A-549 and MCF-7 cells expressing high levels of SOX2 from the three overexpression systems showed higher level expression of these EMT mediators (Fig. 4C and 4D).

**Fig. 4.**
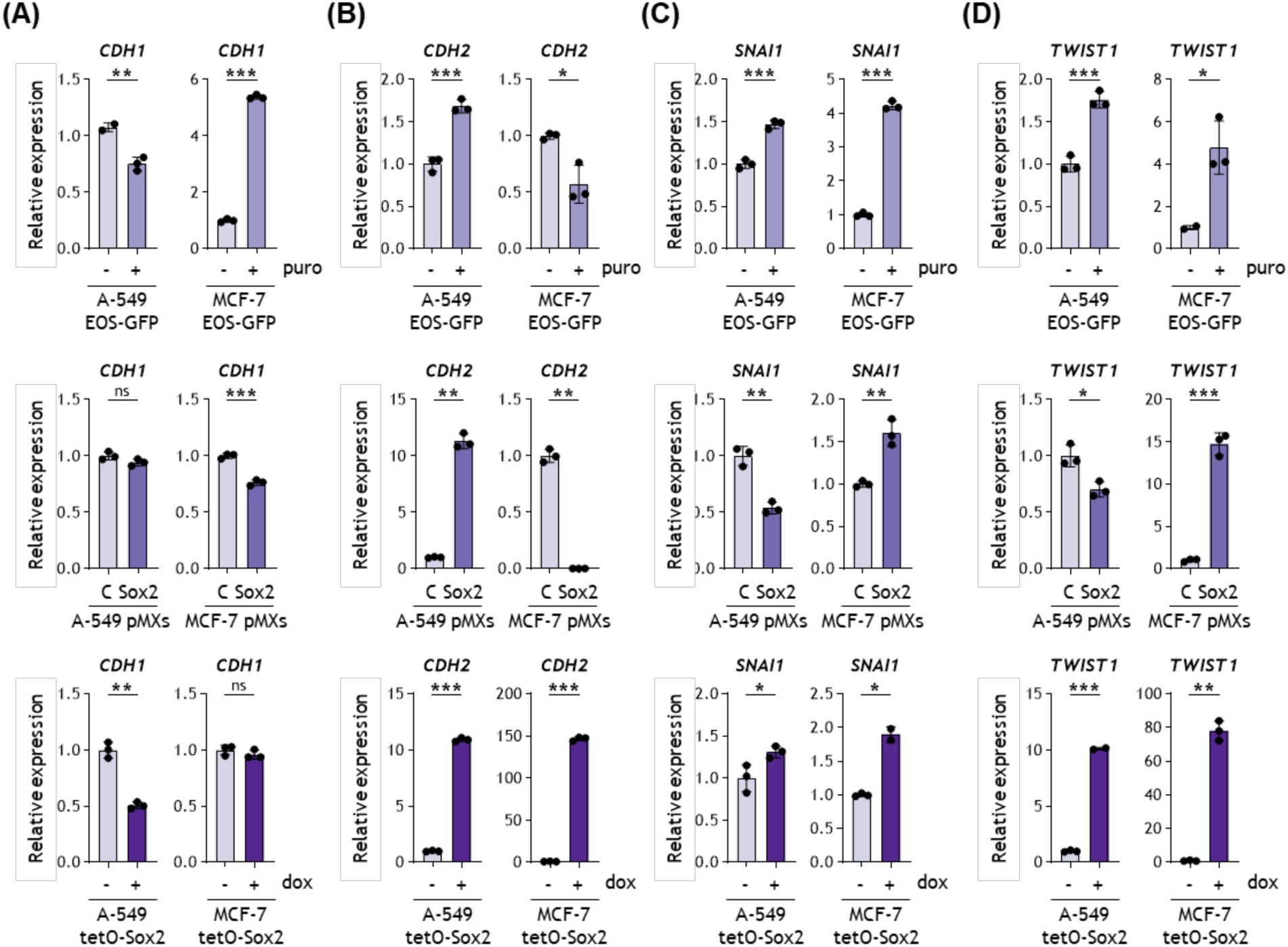
Increasing *SOX2* levels promotes the expression of epithelial-to-mesenchymal transition (EMT) mediators. mRNA expression by RT-qPCR of the epithelial marker *CDH1* **(A)**, the mesenchymal marker *VIM* **(B)** and the EMT effectors SNAI1 and TWIST1 **(C**, **D)** in A-549 and MCF-7 cell lines transduced with EOS-GFP and selected with 1 µg/ml puromycin (top), with pMXs-Sox2-IP or the control pBP-GFP (middle) and pLV-tetO-Sox2 followed by 0.5 µg/ml doxycycline treatment (bottom). puro, puromycin; dox, doxycycline.

Together, the change in cellular morphology, the increased expression of mesenchymal markers and reduced expression of epithelial markers, and the expression of EMT transcriptional regulators, all point to a switch in tumor cell identity that favors a more mesenchymal phenotype.

### Increased levels of SOX2 in cancer cells increase migration

One of the effects derived from the acquisition of an EMT state in cancer cells is the stimulation of cell migration. To test whether this is the case when cancer cells undergo EMT due to increased expression of SOX2, we tested the migration capacity of A-549 and MCF-7 cells forced to express higher levels of SOX2 using our three systems of gradual increased levels. For this, we performed migration assays using transwells. A-549 and MCF-7 cells expressing higher levels of SOX2 were seeded in cell culture medium lacking FBS, and these upper chambers were placed in wells containing medium with 10% FBS to create a chemoattractant gradient that stimulates migration. After incubation, migrating cells were quantified by removing the filter at the bottom of the top compartment, staining, and counting. Again, using the three cellular systems of higher expression of SOX2 we observed a clear and consistent increase in the number of cells migrating in the transwells when SOX2 was overexpressed (Fig. 5A-C).

**Fig. 5.**
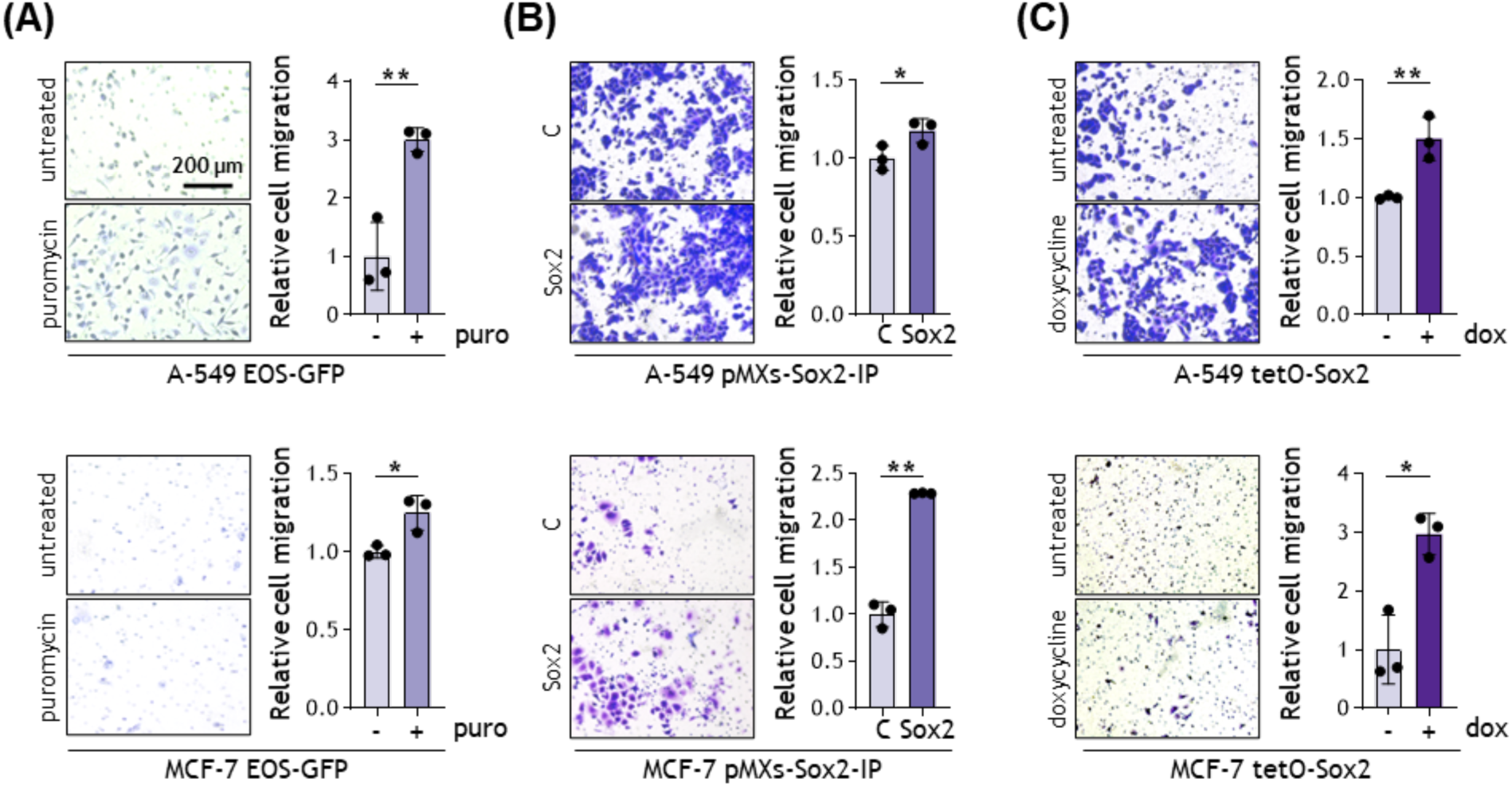
Increasing *SOX2* levels promotes cell migration capacity. Representative images and quantification of the transwell migration assay of A-549 (top) and MCF-7 cells (bottom) transduced with EOS-GFP and selected with 1 µg/ml puromycin **(A)**, with pMXs-Sox2-IP or the control pBP-GFP **(B)** and pLV-tetO-Sox2 followed by 0.5 µg/ml doxycycline treatment **(C)**. C, control; NT, not treated; puro, puromycin; dox, doxycycline.

These assays corroborated the altered cancer cell phenotype observed when SOX2 expression was increased and that are typically observed when epithelial cancer cells increase their plasticity to become more mesenchymal and migratory.

### SOX2 expression is higher in *TP53* mutant human breast cancer patients correlating with a more metastatic phenotype

To further substantiate our molecular findings in cancer patients we decided to analyze the expression of SOX2 in a set of breast cancer patient-derived tissue samples by immunohistochemistry. First, we observed a statistically significant (p=0.036) positive correlation between a TP53+ staining, indicative of a *TP53* mutant status, and a concomitant SOX2+ staining (Table 1). These data reinforce our observations of an inverse correlation between TP53 functionality and *SOX2* expression in cancer cells, with SOX2 protein being expressed at higher levels in tumors with defective TP53. In addition, we observed a clear and statistically significant (p=0.003) difference in SOX2 expression according to the metastatic phenotype of breast tumors, with higher level expression of SOX2 correlating with tumors showing a more metastatic behavior (Table 1).

**Table 1.**
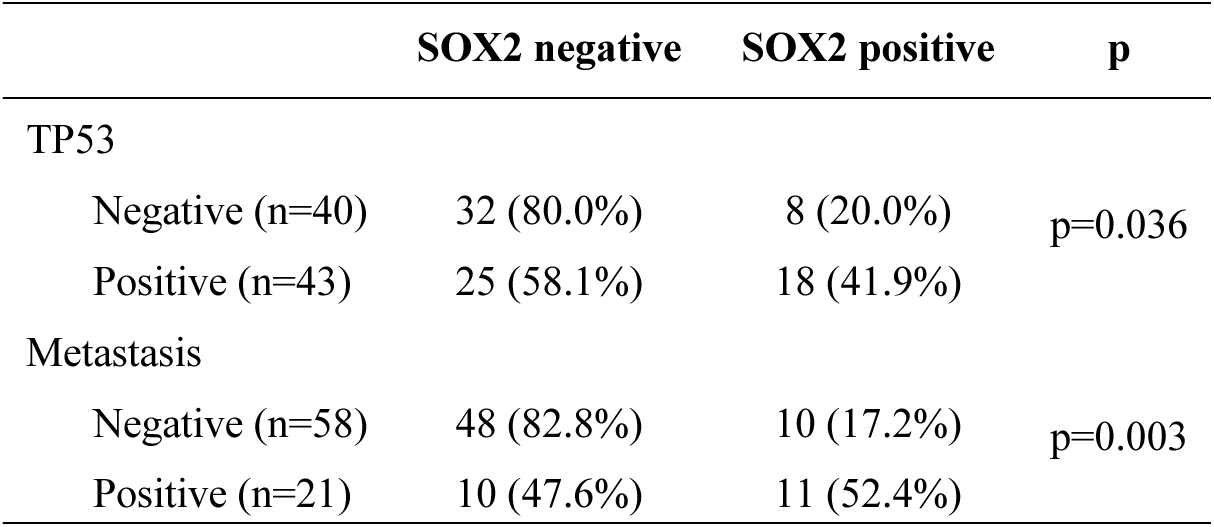
Correlation between SOX2 expression with TP53 status and tumor metastatic state in human breast cancer samples.

|  | **SOX2 negative** | **SOX2 positive** | **p** |
| --- | --- | --- | --- |
| TP53 |  |  |  |
| Negative (n=40) | 32 (80.0%) | 8 (20.0%) | p=0.036 |
| Positive (n=43) | 25 (58.1%) | 18 (41.9%) |  |
| Metastasis |  |  |  |
| Negative (n=58) | 48 (82.8%) | 10 (17.2%) | p=0.003 |
| Positive (n=21) | 10 (47.6%) | 11 (52.4%) |  |

Together, our data in breast cancer patient tissues corroborate our in vitro findings, suggesting that another tumor suppressor function of *TP53* is to restrict SOX2 expression in cancer cells carrying other genetic mutations that could lead to a change in tumor cell identity and the acquisition of a more migratory and invasive activity.

## DISCUSSION

The reactivation of developmental programs is a frequent event during cancer progression that confers tumor cells with advantages in cell competition. *SOX2* is one of the key developmental genes whose expression has been reported to be altered in many tumors. Adult somatic cells typically maintain the *SOX2* locus completely repressed. However, deletion of some key tumor suppressor genes, such as CDK inhibitors *Cdkn1a* and *Cdkn1b*, or the retinoblastoma members, *Rb1* and *Rbl2*, results in a partial de-repression and low-level expression of *Sox2* capable of causing a dramatic phenotype ^10,22^. When these genetic alterations are combined with deletion of *Trp53*, the deregulated expression of *Sox2* is maximal ^11^.

These observations led us to explore the role of TP53 repressing the expression of *SOX2* in cancer. For this, we set up a number of cancer cell lines of different origin and with defective p53 function or retaining a wild-type tumor suppressor protein and genetically reversed the status of p53 on them. This allowed us to robustly prove an inverse correlation between active *TP53* and repressed *SOX2* expression and, conversely, decreased *TP53* and induction of *SOX2* expression (Fig. 1A-D). In the context of pluripotency and the transition to a differentiated state, Trp53 has already been shown to be capable of regulating pluripotency factors ^23,24^. For example, mouse ES cell differentiation caused by the DNA damaging agent doxorubicin causes a shift in whole genome occupancy of Trp53, resulting in its recruitment to regions in which pluripotency core transcription factors reside and showing a repressive activity acting through distal enhancers, while at the same time it binds promoter regions of differentiation-associated genes to activate them ^24^. Interestingly, we have recently described new distal developmental enhancers controlling the expression of Sox2 (namely *SRR124-134*) that can be reprogrammed during tumorigenesis that merit further detailed investigation ^25^.

Our results using luciferase reporter plasmids suggest that this repressive control over *SOX2* expression by TP53 is transcriptionally mediated (Fig. 3A). The ChIP analysis we performed points to the binding of Trp53 to the *Sox2* promoter region and to the enhancer *SRR2* (Fig. 3B). This *SRR2* was shown to be bound by Sox2 together with Oct3/4 to activate its own transcription, and was originally identified together with *SRR1* as regulatory regions stimulating *Sox2* transcription in ES cells and promoting the undifferentiated state ^26^. Interestingly, an activated *SRR2* showing increased Sox2 binding capacity in cancer cells under oxidative stress has been reported to confer resistance to this type of stress, contributing to tumorigenesis ^27^. This link between increased levels of expression of *Sox2* and the activity of the *SRR2* was also reported in triple negative breast cancer cells. A highly responsive *SRR2* is a feature of a highly tumorigenic subpopulation in triple negative breast cancers showing increased capacity to form tumor spheres ^27^. Thus, having a transcriptional repressive factor such as TP53 acting directly at this location could have profound effects on *SOX2* expression in cancer cells. There are, however, other ways to repress *SOX2* expression in a TP53-dependent manner different from a direct effect of TP53 that we have not explored in the context of cancer. For example, microRNA miR-34a has been shown to represent a barrier for somatic cell reprogramming and this suppression of reprogramming was shown to be due, at least in part, to repression of pluripotency genes, including *SOX2* ^28^.

The role of SOX2 in cancer has been addressed by many laboratories around the world and found to be important in the development, maintenance of stemness, cancer progression and resistance towards cancer therapies ^8,9,29^. *SOX2* was originally identified as an oncogene amplified in squamous cell carcinomas of the lung and esophagus where it drives the expression of markers of both squamous differentiation and pluripotency ^6^. Interestingly, in prostate cancer, mutations in *TP53* and *RB1* lead to increased expression of *SOX2* that results in increased cellular plasticity, allowing the escape from antiandrogen therapy by lineage switching. Prostate cells with high levels of *SOX2* shift from an androgen receptor-dependent luminal epithelial phenotype to a basal-like state that is independent of the androgen receptor, making them resistant to the antiandrogen drug enzalutamide ^30,31^. On another report, using a reporter plasmid based on the activation of GFP expression driven by genetic elements bound and activated by SOX2, the authors showed how selecting GFP+ cells from different breast cancer cell lines allows the isolation of a subpopulation with higher tumor- and metastasis-initiating activity, and resistance to chemotherapy ^32^. In our case, using systems that allow the generation of cancer cells showing gradually increased levels of *SOX2* we observed a phenotypic change that morphologically results in an epithelial-to-mesenchymal transition (Fig. S1 and S2). This was further confirmed molecularly by analyzing the levels of epithelial and mesenchymal markers, and by measuring the levels of EMT executers (Fig. 4A-D). All these markers clearly show that cancer cells with high levels of *SOX2* tend to acquire a phenotype that resembles a more mesenchymal state. Since EMT is a process shared by development and cancer that is linked with the capacity to switch cellular states, a process that is related with cell migration capacity ^33^ we decided to test whether *SOX2* expressing cancer cells show an increased capacity to migrate. Our results clearly show that higher levels of *SOX2* in different cancer cells cause an increased capacity to migrate (Fig. 5A-C). These link between altered expression of *SOX2* caused by mutation of *TP53* and a more migratory and metastatic phenotype could also be substantiated with patient derived samples in which we observed a correlation between these parameters, reinforcing our experimental results obtained with manipulated cancer cell lines in vitro (Table 1).

Altogether, our results show that among the range of mutations and genetic alterations occurring to cancer cells, those that impact the regulation of *SOX2* expression and that act in concert with TP53 deficiency, lead to increased expression of this developmental factor, conferring new properties to cancer cells related with a more plastic cell identity and increased migration. A better understanding of the regulation of *SOX2* expression might provide clues for therapeutic interventions aimed at reducing cancer cell plasticity, contributing to an improved management of neoplastic diseases.

## Abbreviations

ChIP: Chromatin Immunoprecipitation
DMEM: Dulbecco’s Modified Eagle Medium
EMT: Epithelial-to-Mesenchymal Transition
FBS: Fetal Bovine Serum
GFP: Green Fluorescent Protein
PEI: Polyethylenimine
PFA: Paraformaldehyde
RT-qPCR: Quantitative Real Time PCR
shRNA: short hairpin RNA
SOX2: *SRY-box transcription factor 2*
SRR1: *Sox2 Regulatory Region 1*
SRR2: *Sox2 Regulatory Region 2*
TMA: Tissue Microarray

## STATEMENTS AND DECLARATIONS

## ACKNOWLEDGEMENTS

We are grateful to José Antonio Costoya (CIMUS-USC, Spain) for the kind gift of T653 and T731 mouse glioma cell lines. The graphical abstract was created with BioRender.com.

## CONSENT TO PARTICIPATE

Not applicable.

## CONSENT FOR PUBLICATION

Not applicable.

## ETHICS APPROVAL

Not applicable.

## AUTHOR CONTRIBUTIONS

P.L.-F. and J.M.V. had an equal experimental and intellectual contribution and are responsible for most of the data; T.F., C.C., S.D.S.-A., V.E.-S., and P.P. helped with specific experiments and data analysis; M.G.-B., L.E.A., J.A.M., and C.R. provided particular reagents, helped with the execution of experiments and their analysis; G.M.-B. provided the human sample study; A.V. and M.C. conceived, designed, directed the project, and analyzed the data; P.L.-F., A.V. and M.C. wrote the manuscript. All authors read and approved the final manuscript.

## DATA AVAILABILITY

All data generated or analyzed during this study are included in this published article. The materials used in this study are available from the corresponding authors, A.V. and M.C., upon reasonable request.

## SUPPLEMENTARY INFORMATION

### MATERIALS AND METHODS

#### Flow cytometry

The percentage of A-549 and MCF-7 EOS-GFP cells that were GFP positive was determined using a FACScalibur Flow Cytometer (BD Biosciences).

### FIGURES AND LEGENDS

**Fig. S1.**
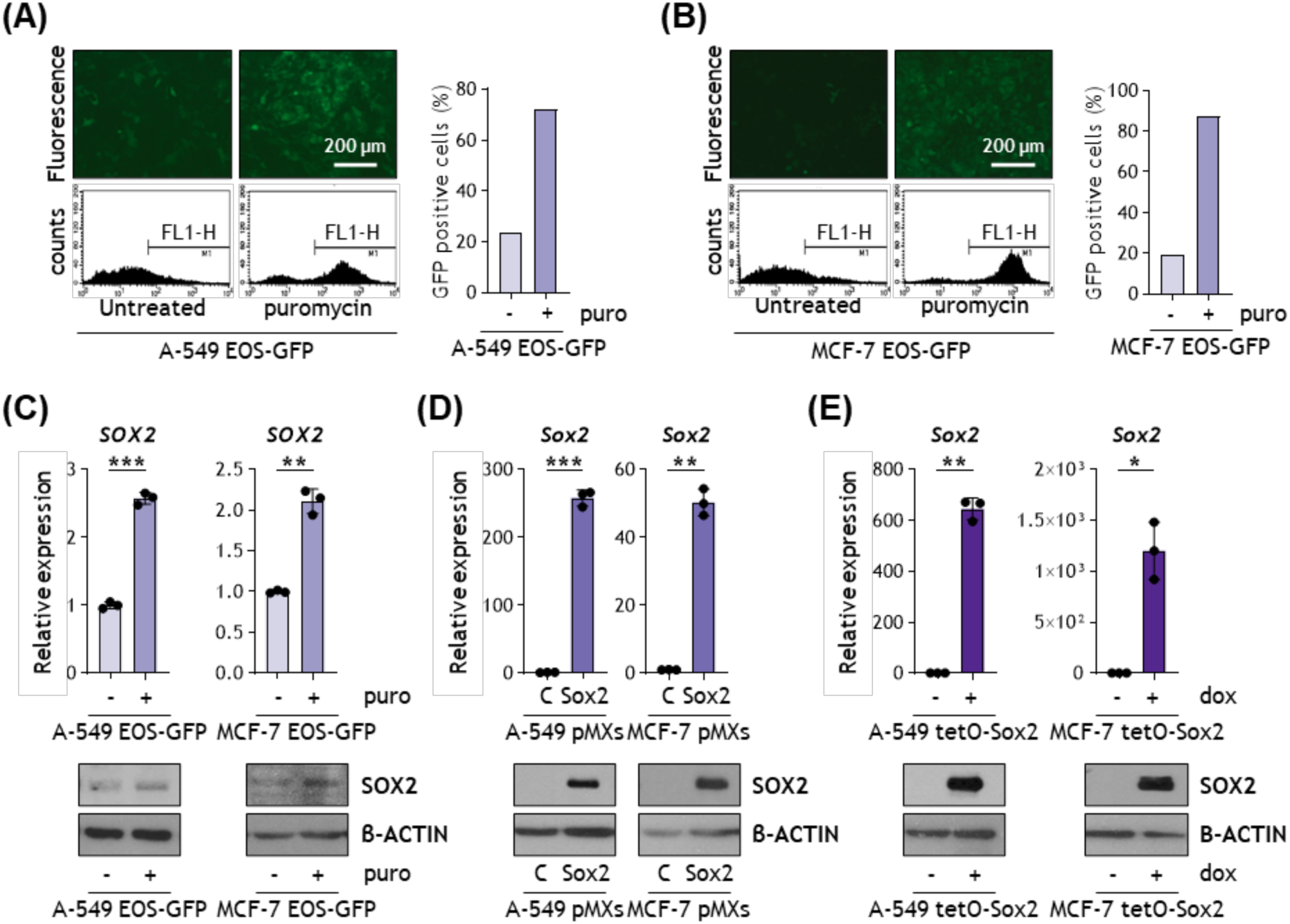
Generation of lung and breast tumor cell lines with increased *SOX2* expression. Representative fluorescence images (top) and flow cytometry graphs (bottom) to detect GFP expression in A-549 **(A)** and MCF-7 cells transduced with EOS-GFP vector and selected with puromycin **(B)**. *SOX2* mRNA expression by RT-qPCR and SOX2 protein levels by western blot in A-549 and MCF-7 cells transduced with EOS-GFP vector and selected with puromycin **(C)**. *Sox2* mRNA expression by RT-qPCR and protein levels by western blot in A-549 and MCF-7 cells transduced with pMXs-Sox2-IP or the control pBP-GFP **(D)** or with pLV-tetO-Sox2 followed by 0.5 µg/ml doxycycline treatment **(E)**. C, control; NT, not treated; puro, puromycin; dox, doxycycline

**Fig. S2.**
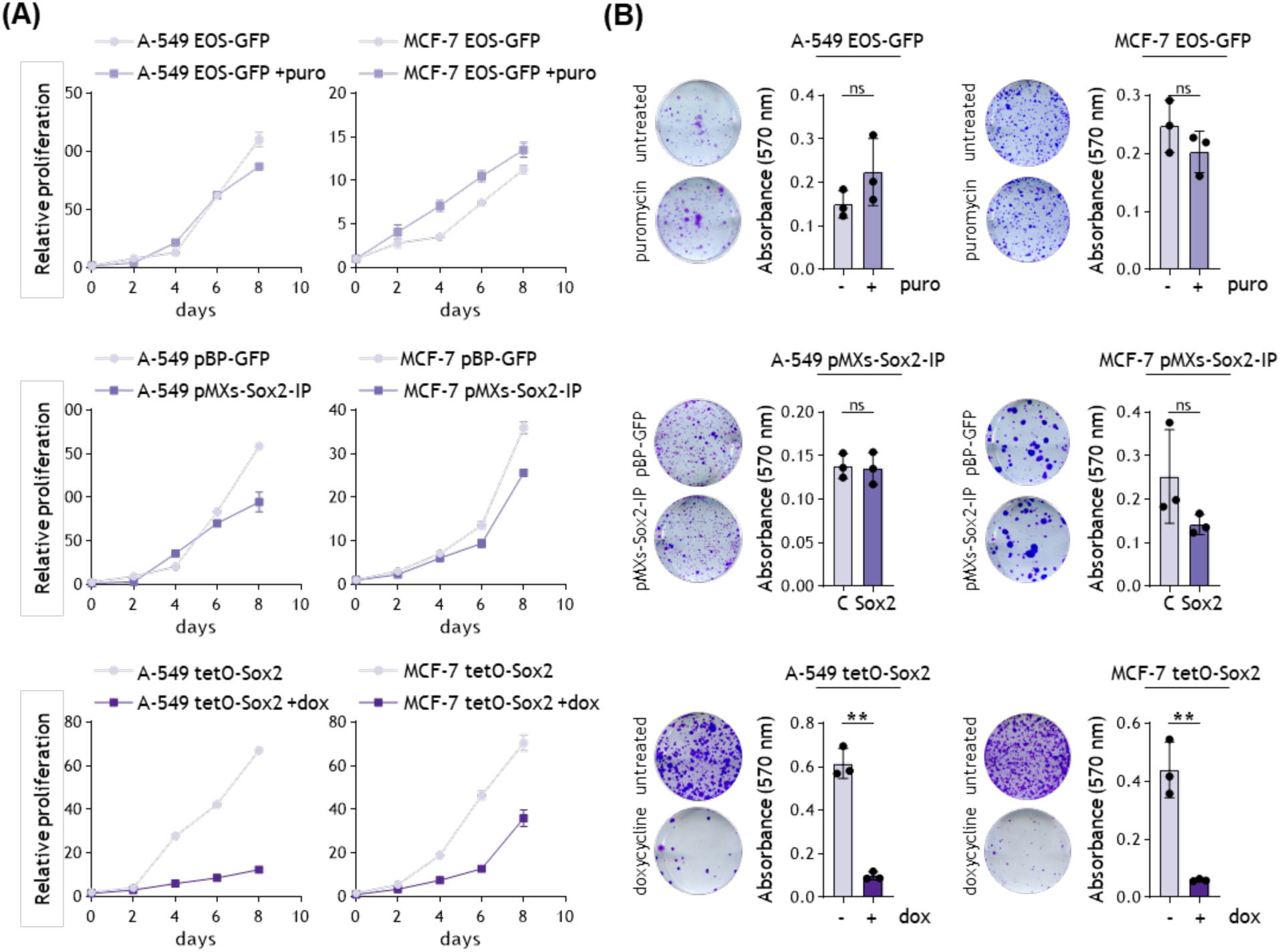
Increasing *SOX2* levels does not increase cell proliferation. **(A)**, Total cell proliferation of A-549 (left) and MCF-7 (right) cell lines transduced with EOS-GFP and selected with 1 µg/ml puromycin (top), with pMXs-Sox2-IP or the control pBP-GFP (middle) and pLV-tetO-Sox2 followed by 0.5 µg/ml doxycycline treatment (bottom). **(B)**, Scans and quantification of the cell proliferation assay of A-549 (left) and MCF-7 (right) cell lines transduced with EOS-GFP and selected with 1 µg/ml puromycin (top), with pMXs-Sox2-IP or the control pBP-GFP (middle) and pLV-tetO-Sox2 followed by 0.5 µg/ml doxycycline treatment (bottom). C, control; NT, not treated; puro, puromycin; dox, doxycycline.

**Fig. S3.**
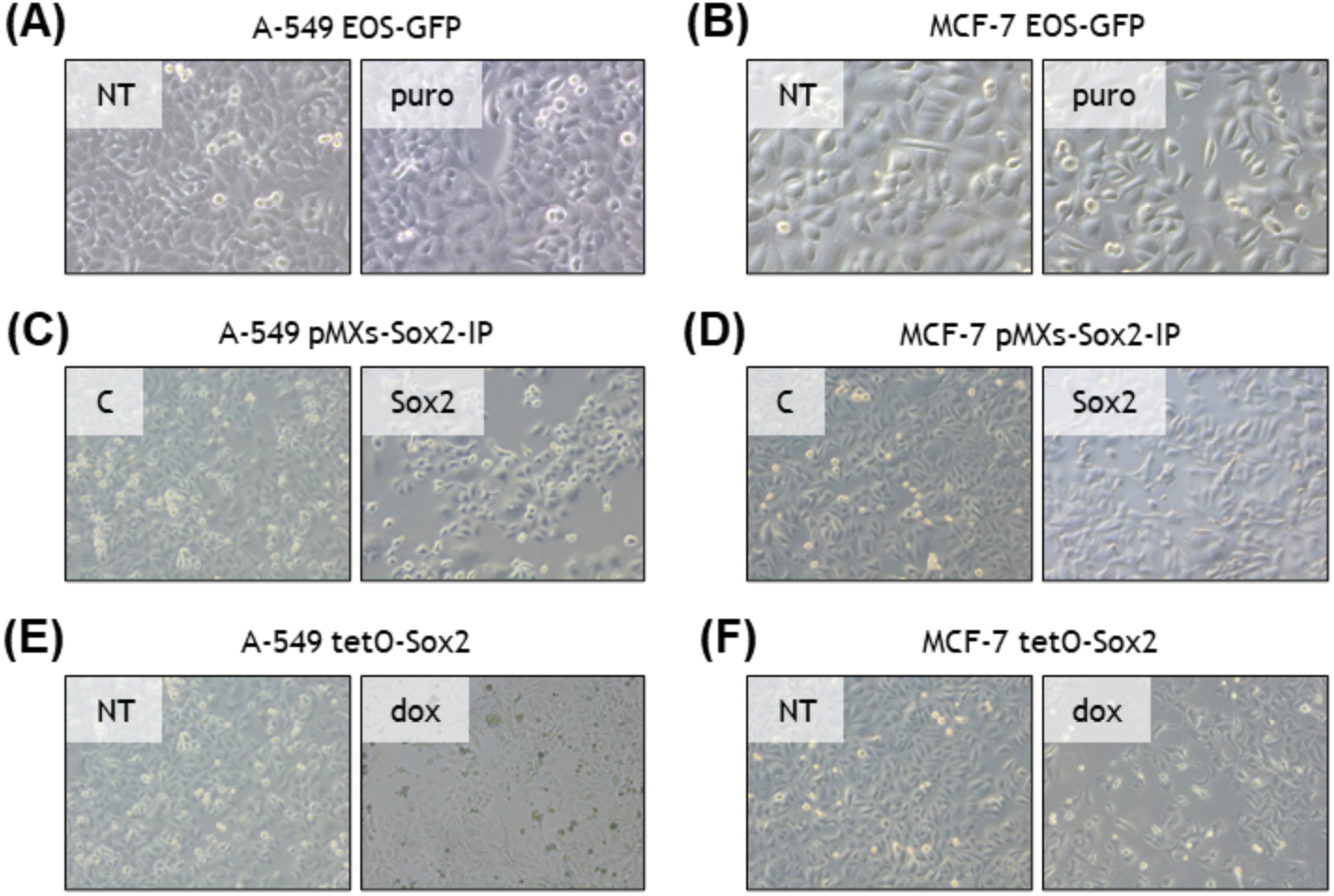
Elevated Sox2 expression causes a morphological change to a mesenchymal phenotype. Representative brightfield images of A-549 and MCF-7 cells transduced with EOS-GFP and selected with puromycin **(A, B)**; transduced with pMXs-Sox2-IP **(C, D)** or with pLV-tetO-Sox2 and induced with doxycycline **(E, F)**. C, control; NT, not treated; puro, puromycin; dox, doxycycline.

